# Tube to Tumour: an integrative epigenomic analysis of DNA methylation in high-grade serous ovarian cancer and precursor serous tubal intraepithelial carcinoma

**DOI:** 10.64898/2026.06.10.731305

**Authors:** Asia Jordan, Aideen McCabe, Alejandra Rodriguez, Kellie Dean, Sudipto Das, Antoinette S. Perry

**Affiliations:** School of Biology and Environmental Science, O’Brien Science Centre, University College Dublin, Ireland; Cancer Biology and Therapeutics laboratory, UCD Conway Institute of Biomolecular and Biomedical Science; Biochemistry and Cell Biology, University College Cork, Cork, Ireland; The Research Centre for Research Training in Genomics Data Science, Galway, Ireland; School of Pharmacy and Biomedical Sciences, Royal College of Surgeons, Dublin, Ireland

**Keywords:** High grade serous ovarian cancer, serous tubal intraepithelial carcinoma, DNA methylation, CpG islands, *cis*-regulatory regions

## Abstract

Serous tubal intraepithelial carcinoma (STIC) is a known precursor of high-grade serous ovarian cancer (HGSOC). Yet, molecular events driving progression from STIC to HGSOC remain poorly defined. Aberrant DNA methylation is a hallmark of cancer, yet its role in early HGSOC remains unclear. We performed a comprehensive meta-analysis of publicly available Illumina Infinium DNA methylation EPIC array datasets assessing 255 samples comprising STIC, HGSOC, and histologically normal fallopian tube tissues. We mapped DNA methylation alterations during early tumorigenesis, identified conserved methylation patterns across STIC and HGSOC, and assessed RNA-sequencing data to define transcriptional consequences within genomic and epigenomic landscapes. STIC and HGSOC exhibited widespread DNA hypomethylation relative to normal tissue, accompanied by focal hypermethylation in CpG islands and 5′ regulatory regions. DNA Hypomethylation intensifies during progression from STIC to HGSOC, particularly in *cis*-regulatory enhancer domains and intergenic regions. We identified 11,660 CpG sites and 447 genomic regions with conserved DNA methylation patterns across STIC and HGSOC. Within these, 70 genes showed coordinated DNA methylation and expression changes, including *TRIM15*, *NKAPL*, and *RIPPLY3*. These findings reveal that epigenetic remodelling occurs in STIC lesions, prior to malignant transformation. DNA methylation alterations at regulatory regions may drive invasion and offer novel avenues for early detection and targeted intervention.

## Introduction

Ovarian cancer is the most lethal gynaecological malignancy, predominantly affecting post-menopausal women. Outcome is strongly tied to disease stage at diagnosis, with 5-year survival rates of 92%, 72%, and 31% when patients are diagnosed at localized, regional and distant stages respectively^1^. In western countries, type 2 ovarian cancers, specifically the high-grade serous ovarian cancer (HGSOC) subtype, are the most common. Type 2 ovarian cancers are more aggressive than their type 1 counterparts^2^. HGSOC is typically diagnosed at a late-stage and subsequently has very poor survival^3^.

Over the last decade, it has emerged that many HGSOCs arise from serous tubal intraepithelial carcinoma (STIC) lesions within the fallopian tube in addition to the ovarian surface epithelium^4^. This delayed understanding of the dual origin has led to challenges in deciphering the carcinogenesis of HGSOC. Signature HGSOC DNA mutations (*TP53*, *BRCA1*, *BRCA2* and *PTEN*) have been detected in STICs^5^. In addition, adjacent histologically normal fallopian tube occasionally harbours similar *TP53* mutations, known as p53 signature lesions, which are believed to progress into STICs, and subsequently HGSOC^5^.

It is estimated that STICs progress to ovarian cancer in ∼6.5 years, with some cases progressing more rapidly (<2 years)^5^. Identifying the molecular characteristics of STIC lesions, which drive disease progression could provide a much-needed insight into the early stages of HGSOC oncogenesis. Early-stage HGSOC tissue is rare due to late-stage diagnosis and is difficult to acquire for research purposes^3^. While STIC lesions may bridge this knowledge gap, they also present their own challenges, namely their small size (<1-11mm) and macroscopic invisibility^6^. The difficulty associated with their collection, in addition to the delayed recognition of their contribution to HGSOC development has limited our understanding of the earliest stages of this disease.

Aberrant DNA methylation is a key characteristic of cancer and broadening our understanding of its role has opened new avenues for improved diagnostics and treatment^7^. Epithelial ovarian cancer displays genome-wide DNA hypomethylation, with hypermethylation mostly occurring at specific CpG islands^8^. Recently, it has been reported that DNA hypermethylation alterations can arise in STIC and are maintained or progressively alter as disease evolves into HSGOC^9^. This new discovery raises several important unanswered questions: what is the role, if any, of DNA hypomethylation in STIC? What is the epigenomic and genomic landscape of DNA methylation alterations in STICs and how does this differ from HGSOC? What is the functional role of DNA methylation alterations in HGSOC? The aim of this study was to address this knowledge gap through a robust meta-analysis of publicly available DNA methylation data to better understand the methylation landscape of STIC and HGSOC.

## Methods

### Data selection

Raw Illumina Infinium DNA methylation EPIC array datasets: GSE133556^10, 11^, GSE155311, GSE155760^9^, GSE168930^8^ and GSE211686^12, 13^ were sourced via the NCBI gene expression omnibus (GEO)^14^. From these datasets we gathered data from primary HGSOC tumours (n=207), STIC precursor lesions (n=11), dormant STIC (dSTIC) (n=1), a p53 lesion (n=1) and histologically normal fallopian tube (hnFT) controls (n=35) (**Supplementary Table 1)**. The total cohort amounted to 255 tissue samples.

These samples were divided into three groups: a matched cohort, which contained paired samples from the same patients (STIC: n=11, dSTIC: n=1, p53 lesion: n=1 and hnFT; n=12), an unmatched cohort (HGSOC: n=140, hnFT: n=23 and STIC: n=11), and a verification cohort (HGSOC: n=65). HGSOC samples were randomly assigned to the unmatched and verification cohorts (**Figure 1**).

**Figure 1.**
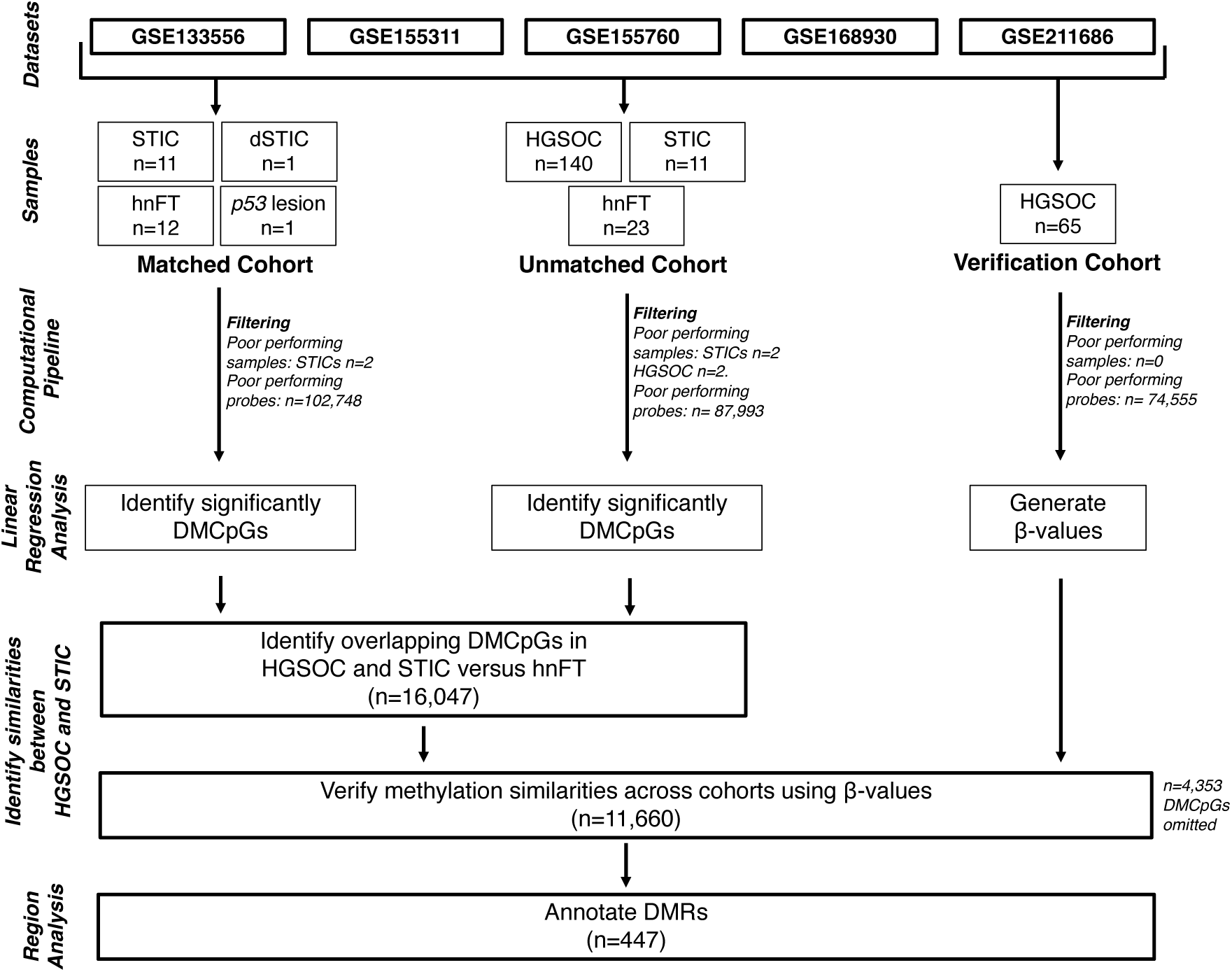
Analysis flowchart. Abbreviations: (HGSOC) High-grade serous ovarian cancer, (STIC) serous tubal intraepithelial carcinoma, (dSTIC) dormant STIC, (hnFT) histologically normal fallopian tube, (DMCpGs) statistically significant differentially methylated CpG sites, (DMRs) statistically significant differentially methylated regions. hnFT samples (n=12) were matched to STIC (n=11), dSTIC (n=1), and p53 (n=1) lesions from the same patients in the matched cohort. The same STIC samples were used across the matched and unmatched cohorts, due to limited available of STIC DNA methylation data.

### Computational Analysis

Raw illumina Infinium DNA methylation EPIC array datasets were processed using the *RnBeads* computational pipeline and annotated to hg19/GRCh37 human genome assembly^15^. Data were normalised via SWAN^16^. Probes and samples were filtered via the greedycut algorithm to remove cross-reactive, poor performing probes that covered SNPs within their last 5 bases or were non-CpG probes (**Supplementary Table 2).** Batch correction was conducted between datasets via surrogate variable analysis using the COMBAT method^17^. We then conducted differential methylation analysis of CpG sites via linear regression. We quantified the total number of hypermethylated and hypomethylated CpG sites for each group comparison using summarized methylation counts. A chi-squared test was applied to assess significance across comparisons.

### Identification of concordantly differentially methylated CpG sites in STIC and HGSOC

Differential DNA methylation analysis was conducted between HGSOC vs hnFT, STIC vs hnFT (both unmatched analyses) and STIC vs hnFT (matched cohort). Statistically significant differentially methylated CpG sites (DMCpGs) had a false discovery rate (FDR) p-value <0.05 and minimum mean β-value difference of ≥20% (0.2) between either HGSOC or STIC compared to hnFT. DMCpGs present in all three comparisons (n=16,047), were brought forward for further analysis (**Supplementary Figure 1)**.

We then verified that the β-values of the DMCpGs were similar between samples of the same type across the different cohorts. We omitted DMCpGs that had ≥10% (≥0.1) difference in their mean β-value between like tissues. The remaining overlapping DMCpGs (overlapping DMCpGs) were retained for the analysis of differentially methylated regions (DMRs). Lastly, we examined whether these overlapping DMCpGs were hyper/hypomethylated concordantly in both STICs and HGSOC when compared to hnFT via unsupervised hierarchical clustering using Euclidian distancing, via complexheatmaps (version 2.22.0)^18^.

### Adjusting for probe distribution on Infinium DNA methylation EPIC array

Probes were annotated to CpG sites using the Illumina Infinium DNA methylation EPIC array manifest (MethylationEPIC_v-1-0_B2)^19^. Each DMCpG site was categorized according to four annotation features from the manifest file: 1) chromosome, 2) epigenetic (e.g. CpG islands, shores and shelves), 3) genetic (e.g. transcriptional start sites, gene body or non-coding intergenic regions according to the UCSC annotation) and 4) *cis-*regulatory elements (e.g. enhancers, determined via Fantom4 and Fantom5). As the EPIC probe distribution is non-random, we calculated an adjusted “relative distribution” (%) by determining the ratio of all probes within each domain (e.g., CpG island, shore, shelf, open sea) on the array and then dividing the number of DMCpGs (hyper/or hypomethylated) by the ratio calculated. This adjustment allows for direct comparison of DNA methylation patterns across genomic and epigenetic domains independent of probe design bias on the EPIC array.

### Annotation of differentially methylated sites and regions

Overlapping sites were input into DMR analysis, which was conducted using the DMRcate^20^ and limma^21^ packages. Linear regression was used to assess DMRs between HGSOC and hnFT. Regions were required to have ≥3 DMCpGs and considered differentially methylated if they had a mean β-value difference of ≥10% (0.1), Stouffers p-value <0.05, multiple comparison statistic and harmonic mean FDR p value <0.05. Regions were also annotated using the Illumina Infinium DNA methylation EPIC array manifest (MethylationEPIC_v-1-0_B2)^19^.

### Correlations between DMRs and RNA-sequencing data

The expression of genes associated with each DMR was assessed using two RNA-sequencing (RNA-seq) datasets obtained from the European Nucleotide Archive (ENA), containing raw FASTQ reads for HGSOC (n=25) and hnFT (n=27) (PRJNA926127, PRJNA398141)^22^. No RNA-seq datasets were available for STIC samples.

Raw FASTQ reads were obtained from the ENA-Browser, and quality control checks were performed using the FastQC package^23^. Raw FASTQ files were trimmed and quality-checked using Trim Galore v0.6.10^24^ with a quality threshold of 20, a minimum length of 75bp and stringency 3 for adapter trimming. Transcript level abundances were generated from trimmed RNA-seq reads using Salmon with the GRCh37 (hg19) SAF index, employing bias correction for GC content and sequencing biases, and alignment-free verification^25^. A gene level matrix was generated via the *tximport* package using R (version 4.2.2), which was then used to construct a DESeqDataSet object^26^. Differential expression analysis was conducted to identify genes deregulated between HGSOC and hnFT samples, accounting for batch as a covariate (Love, Huber and Anders, 2014). A log2FoldChange of >1 and <-1 and an adjusted p-value threshold of 0.05 were applied to identify differentially expressed genes.

## Results

Identifying DNA methylation alterations that occur in STIC, specifically those that are maintained throughout HGSOC progression, may offer key insights into disease onset and carcinogenesis. Epigenetic aberrations may be enablers of the metastatic potential in STIC by altering the transcriptomic and subsequent proteomic profiles of each cell. In addition, understanding these molecular alterations may provide clinical opportunities for early-stage diagnosis, subsequently improving treatment outcomes.

### HGSOC and STIC are differentially methylated and predominantly hypomethylated relative to hnFT

We first examined the overall DNA methylation patterns in hnFT, STIC and HGSOC via principal component analysis (PCA), and found that the biggest driver in DNA methylation variance was tissue pathology (**Supplementary Figure 2)**. Notably DNA methylation of the p53 lesion clustered with hnFT tissues, while the singular dSTIC lesion was clustered with other STICs, and STICs seemed indistinguishable from HGSOC. Batch effect differences were assessed through PCA and SVA, and no surrogate variable were identified.

Analysis of differential methylation between unmatched HGSOC to hnFT and STIC to hnFT yielded similar quantities of DMCpGs: n=101,995 and n=100,138, respectively (**Table 1**). Methylation differences between matched STIC lesions and hnFT produced notably fewer DMCpGs (n=29,159) suggesting that some differences observed in the unmatched analysis may be interindividual. The key finding across each of the four comparisons was the predominance of DNA hypomethylation in both HGSOC and STIC. The frequency of DNA hypermethylation in STIC (34.15%) was more than double that of HGSOC (14.46%), when compared to hnFT (p ≤ 0.0001, chi squared test of independence). These results suggest progressive shifts in methylation patterns from normal healthy tissue (hnFT) to precursor lesions (STIC) and invasive carcinoma (HGSOC).

**Table 1.**
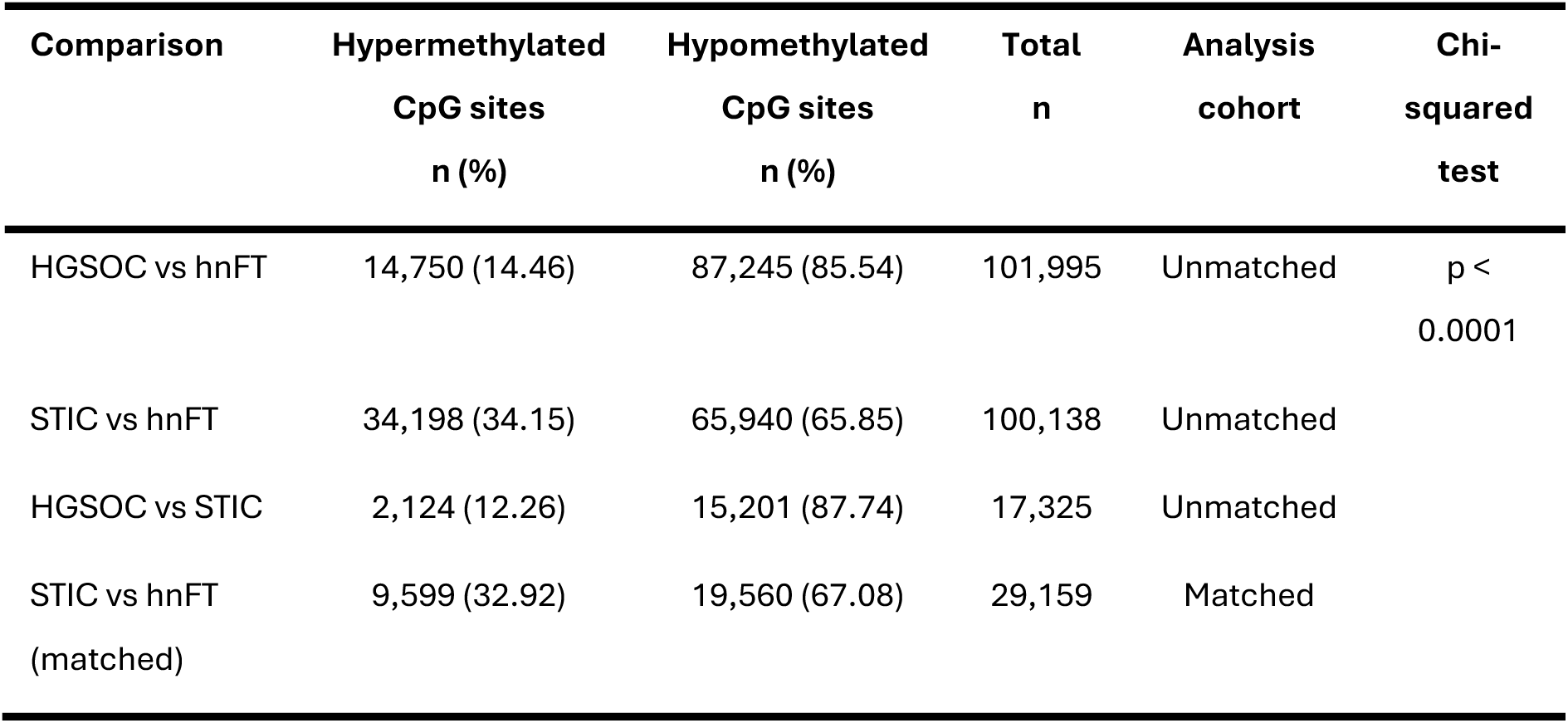
Significantly differentially methylated CpG sites and regions in STIC and HGSOC.

Comparing the DNA methylation states of HGSOC and STIC yielded a far smaller number of DMCpGs (17,325) demonstrating a stronger similarity between these tissues compared to hnFT. The majority of these (87.74%) were again hypomethylated in HGSOC compared to STIC. However, of the top 1000 most significantly DMCpGs, 706 were hypermethylated in the tumours indicating a small yet important role for these hypermethylated sites.

### The genomic and epigenomic landscape of DNA methylation in STIC and HGSOC

We next examined the genomic distribution of the differential methylation in STIC and HGSOC, which may provide insights into potential functional consequences. Looking first at the epigenetic distribution of the DMCpGs, we observed a similar distribution in both STIC and HGSOC compared to hnFT. Notably for the comparisons of STIC to hnFT, regardless of whether the comparison was matched/unmatched, distributions were highly similar (**Figure 2, Supplementary Figure 3)**. For all distributions a chi-squared test of independence confirmed that methylation alterations were not uniformly distributed across the genome. Hypomethylation was predominantly observed at CpG shelves and open sea, whereas hypermethylation showed a more even distribution across islands, shores, shelves and open sea (**Figure 2A**). Similarly, consistency was seen in the genomic distribution of DMCpGs in STIC and HGSOC where the majority (>75%) of hypo and hypermethylation occurred outside of the 5’ regulatory regions (**Figure 2B**). *Cis*-regulatory enhancer domains exhibited significant DNA methylation alterations, showing predominantly hypermethylation in HGSOC and STIC (**Figure 2C**).

**Figure 2.**
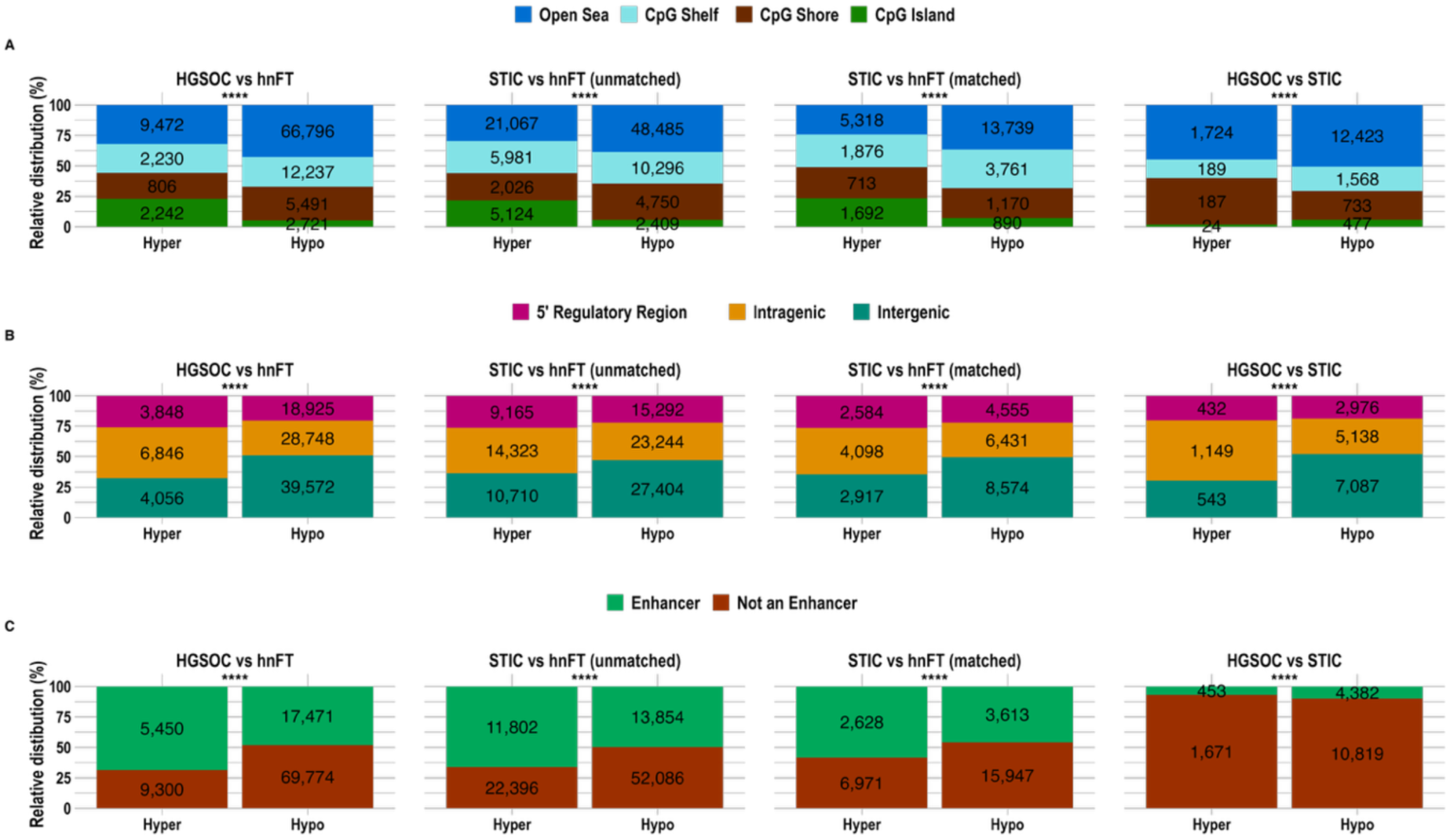
HGSOC and STIC DMCpGs annotation. The relative distribution (%) of hypermethylated (Hyper) and hypomethylated (Hypo) DMCpGs in HGSOC or STIC relative to hnFT (both matched and unmatched) and between HGSOC and STIC across (**A**) epigenetic domains (**B**) genomic domains and (**C**) *cis*-regulatory enhancers. Asterisks indicate statistical significance (**** p < 0.0001) determined using a Chi-squared test of independence comparing proportions of hypermethylated and hypomethylated sites within each domain. Absolute numbers (unnormalized) of DMCpGs are shown for each element.

The ratio of hypermethylated to hypomethylated sites averaged approximately 1:2 in STIC and 1:7 in HGSOC, indicating a progressive loss of DNA methylation with disease advancement. However, certain chromosomes, notably 18 and X, displayed markedly greater hypomethylation, with ratios of 1:5 and 1:10 in STIC, and 1:10 and 1:28 in HGSOC, respectively (**Supplementary Figure 3A-C**). The X chromosome showed a pronounced, and statistically significant progressive reduction in DNA methylation from STIC to HGSOC (p-value < 0.0001, Chi-squared test). While most methylation alterations in STIC lesions occurred on chromosome 18, HGSOC tissues exhibited the greatest number of aberrations on chromosome 8, followed by chromosome 18. These findings suggest that DNA hypomethylation may occur in a chromosome-specific pattern, becoming more extensive as lesions progress from STIC to invasive HGSOC.

Finally, we examined the epigenomic, genomic and chromosomal distribution of differential methylation between STIC and HGSOC. Difference’s in DNA methylation predominantly occurred outside of CpG islands, 5′ regulatory regions, and enhancer domains, suggesting that the epigenetic states of these regions are largely conserved between STIC and HGSOC (**Figure 2)**. Chromosome X exhibited the fewest differentially methylated sites and chromosome 4 the greatest number, when comparing HGSOC to STIC tissues (**Supplementary Figure 3D**).

### Differential DNA methylation patterns associated with HGSOC are established within precursor STIC lesions

Given that STIC lesions are recognized precursors of HGSOC, we investigated whether HGSOC-associated DNA methylation alterations are present at the STIC stage. We identified 16,047 CpG sites that were consistently differentially methylated across HGSOC vs hnFT, STIC vs hnFT (matched) and STIC vs hnFT (unmatched) comparisons. This number reduced to 11,660, when we omitted DMCpGs that had ≥10% (≥0.1) difference in their mean β-value between like tissues (**Supplementary Figure 1**). These shared sites, hereafter referred to as “overlapping DMCpGs,” represent methylation changes common to both STIC and HGSOC relative to hnFT.

We also sought to ensure that the overlapping DMCpGs maintained consistent direction of β-value differences in HGSOC and STIC compared to hnFT, in terms of either hypo– or hypermethylation (**Figure 3A)**, Notably, we observed stronger hypomethylation (lower beta values) in HGSOC and stronger hypermethylation (higher beta values) in STIC relative to hnFT. Consistent with the earlier results, most of the overlapping DMCpGs (9,894) were hypomethylated with 1,766 hypermethylated.

**Figure 3.**
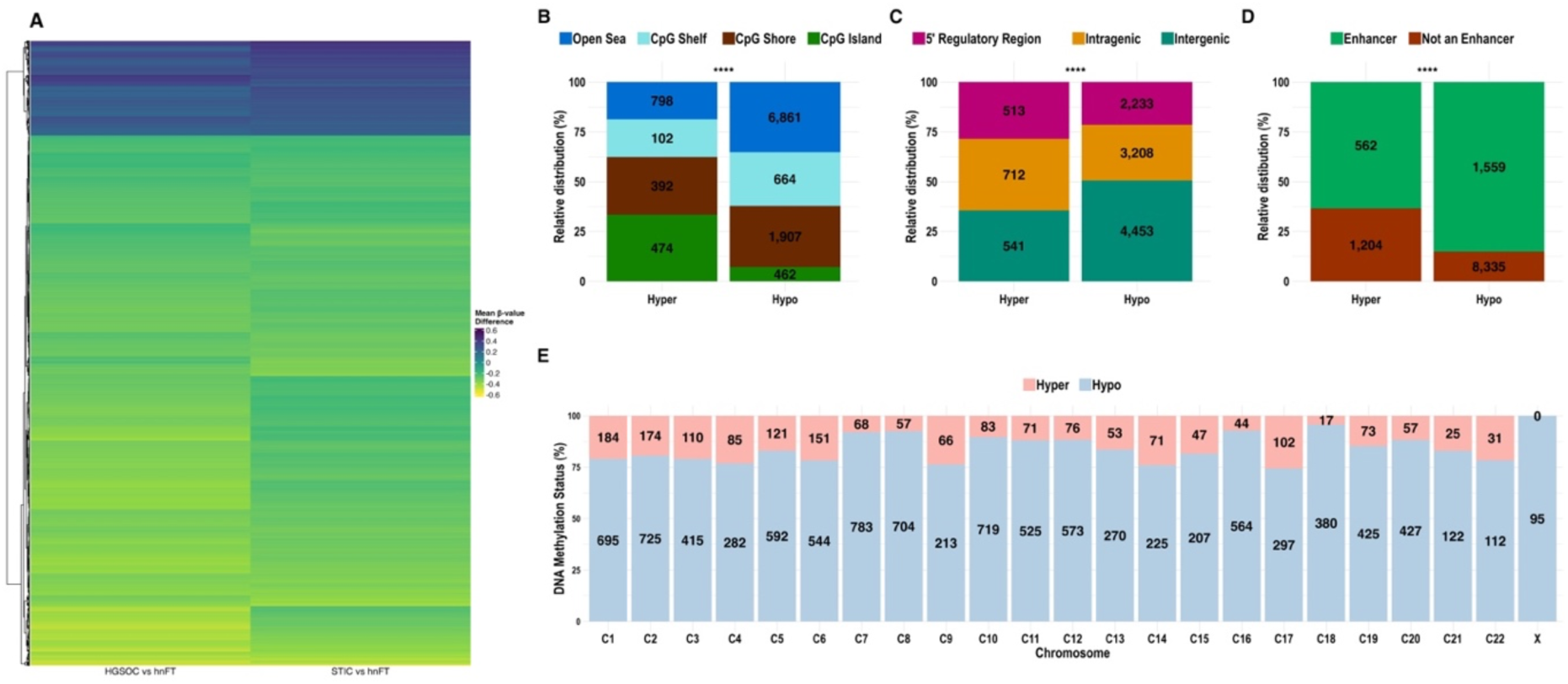
DNA methylation and annotation and of overlapping DMCpGs in HGSOC and STIC. (**A**) Hierarchical clustering using euclidean distancing of the mean β-value differences derived from HGSOC vs hnFT and STIC vs hnFT comparisons A heatmap of the 11,660 overlapping DMCpGs depicting the mean beta value difference between HGSOC and STIC relative to hnFT. The relative distribution of hypermethylated (Hyper) and hypomethylated (Hypo) overlapping DMCpGs across (**B**) epigenetic domains (**C**) genomic domains and (**D**) Enhancers. (**E**) The methylation state of each CpG site is indicated across each chromosome. Asterisks indicate statistical significance (**** p < 0.0001) determined using a Chi-squared test of independence comparing proportions of hypermethylated and hypomethylated sites within each domain. Absolute numbers (unnormalized) of DMCpGs are shown for each element.

We next assessed the distribution of these overlapping DMCpGs to determine whether early methylation alterations exhibit similar regional biases to those observed in HGSOC. The disproportionate methylation patterns observed across epigenomic and genomic elements were also evident within these overlapping DMCpG sites. Epigenetically, these sites showed pronounced hypermethylation within CpG islands (33%), whereas most hypomethylation occurred in CpG shores, shelves, and open sea regions (cumulatively 93%), with only 7% of hypomethylated sites located within CpG islands (**Figure 3B**). Similarly, hypomethylation occurred disproportionately within intergenic and intragenic regions (cumulatively 79%) (**Figure 3C**). Interestingly, many of these overlapping DMCpGs annotated to enhancer regions, 63% of the hypermethylated and 85% of the hypomethylated sites (**Figure 3D**). This suggests the initiation of a potential deactivation of some enhancer domains in STICs during the early stages of disease that persists through HGSOC progression. However, without an understanding for transcription factor binding at these regions it is difficult to conclude whether the increase in hypomethylation is consistent with increased activity. Consistent with these observations, the chromosomal distribution of methylation alterations seen previously (**Supplementary Figure 3D**) was maintained within the overlapping DMCpGs (**Figure 3E**). Chromosome 18 exhibited the greatest degree of consistent differential methylation across both STIC and HGSOC, whereas the X chromosome displayed the least, with no hypermethylated CpG sites conserved between the two tissue types.

### Differentially hypermethylated regions in HGSOC and STIC predominantly occur within regulatory sequences

CpG dinucleotides may impact gene function, depending on their genomic location and their proximity to neighbouring CpG sites. Therefore, we reduced the overlapping DMCpGs to regions (DMRs) and analysed their proximity to regulatory domains. The 11,660 overlapping DMCpGs annotated to 447 DMRs. In keeping with the overriding trend of DNA hypomethylation in STICs and HGSOC, 76.51% (n=342) of the regions were also hypomethylated (**Supplementary Table 3**). Most hypermethylated regions annotated to regulatory elements, namely 5’regulatory regions (49.52%), CpG islands (62.86%) and enhancers (49.52%). However, the opposite was found for hypomethylated regions, which mostly annotated to intergenic regions (39.47%), open sea (50%) and regions of the genome not associated with enhancers (86.55%). Enhancers tended to be hypomethylated and were mostly found at CpG shelves, shores and open sea, whereas the hypermethylated enhancers were enriched at CpG islands (**Figure 4A**). Most hypermethylated regions annotated to 5’ regulatory domains, of which the majority contained CpG islands (58%). In contrast, hypomethylated 5’ regulatory DMRs typically coincided with CpG shores, shelves and open sea (92%). Similarly, intergenic hypomethylated DMRs mostly annotated to open sea (**Figure 4B**).

**Figure 4.**
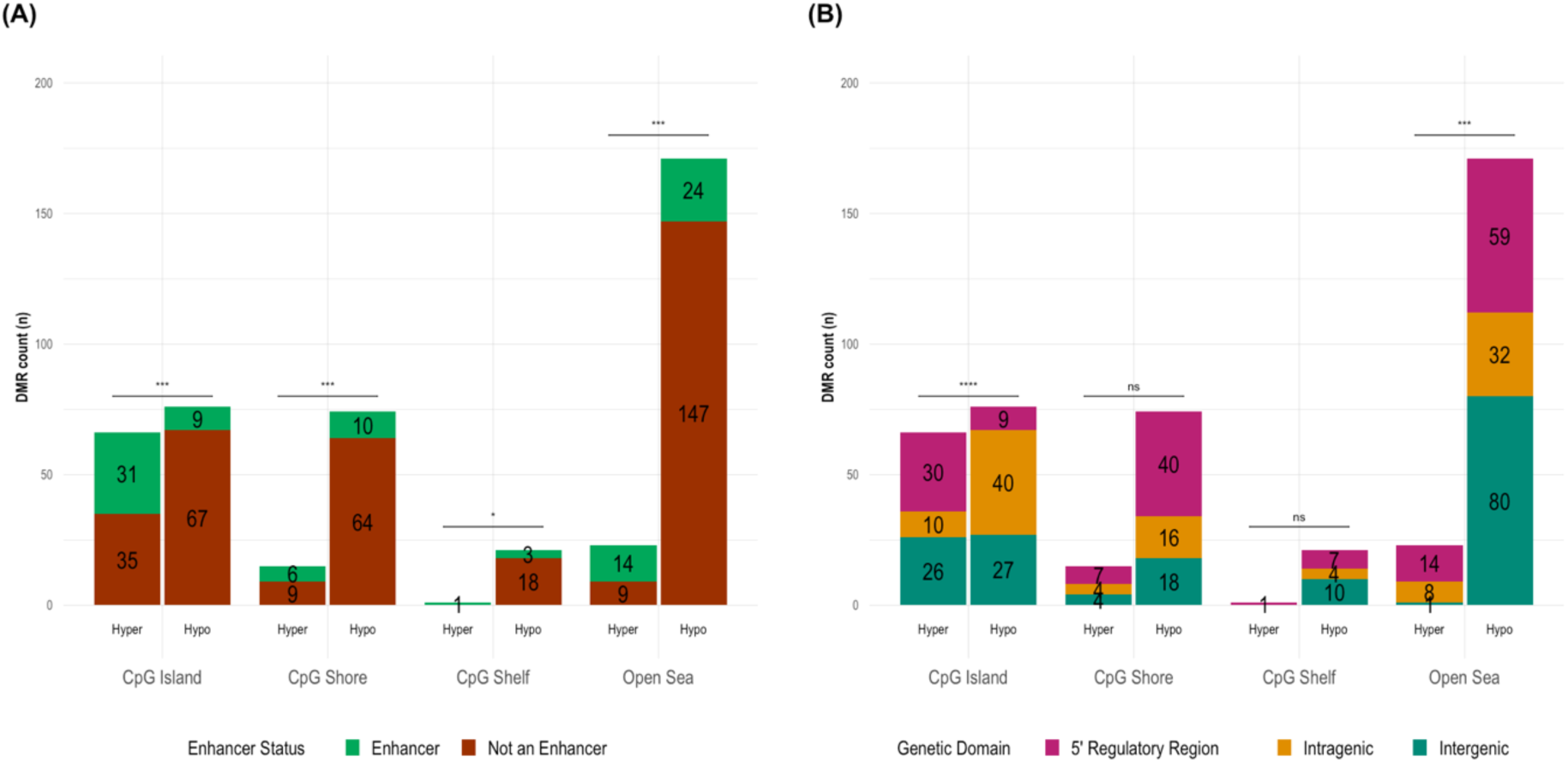
Annotation and β-value distribution of significantly differentially methylated regions (DMRs) in STICs and HGSOC. The annotation of hypermethylation (hyper) and hypomethylated (hypo) DMRs across epigenetic domains (x-axis), stratified by (**A**) enhancer domains and (**B**) genetic domain. Asterisks indicate statistical significance (\*\*\*\**p* < 0.0001, \*\*\**p* < 0.001, ** *p* < 0.01, * *p* < 0.05, ns = not significant.) determined using a Chi-squared test of independence comparing proportions of hypermethylated and hypomethylated sites within each domain.

### Integration of DNA methylation and RNA-seq data identifies differentially expressed and methylated genes in HGSOC

It is widely recognised that abnormal DNA methylation of genes and regulatory elements influences gene expression^27^. We aimed to identify methylation sensitive genes located within DMRs that were also differentially expressed using an independent dataset of HGSOC and hnFT samples. To that end, two independent RNA-sequencing datasets containing n=24 HGSOC samples and n=27 hnFT samples were used as input for the differential expression analysis^22^. Using a log2fold change cut-off of >1 and <-1, and an adjusted p-value threshold of 0.05, we identified 5,966 differentially expressed genes (DEGs) between HGSOC and hnFT, of which 3,168 were upregulated and 2,798 were downregulated **(Supplementary Figure 4).**

Next, the integrative analysis of DNA methylation and gene expression was performed by determining the intersection between DEGs and DMRs. Among the 447 HGSOC/STIC DMRs, 244 overlapped with at least one gene. Some genes were associated with multiple DMRs. Out of 244 DMRs, 222 unique genes were subsequently analysed. Of these 222 genes, 70 were also differentially expressed **(Supplementary Table 4).** These genes were classified into four groups based on their expression and methylation patterns: hypomethylated and upregulated, hypermethylated and upregulated, hypomethylated and downregulated, and hypermethylated and downregulated (**Figure 5A**).

**Figure 5:**
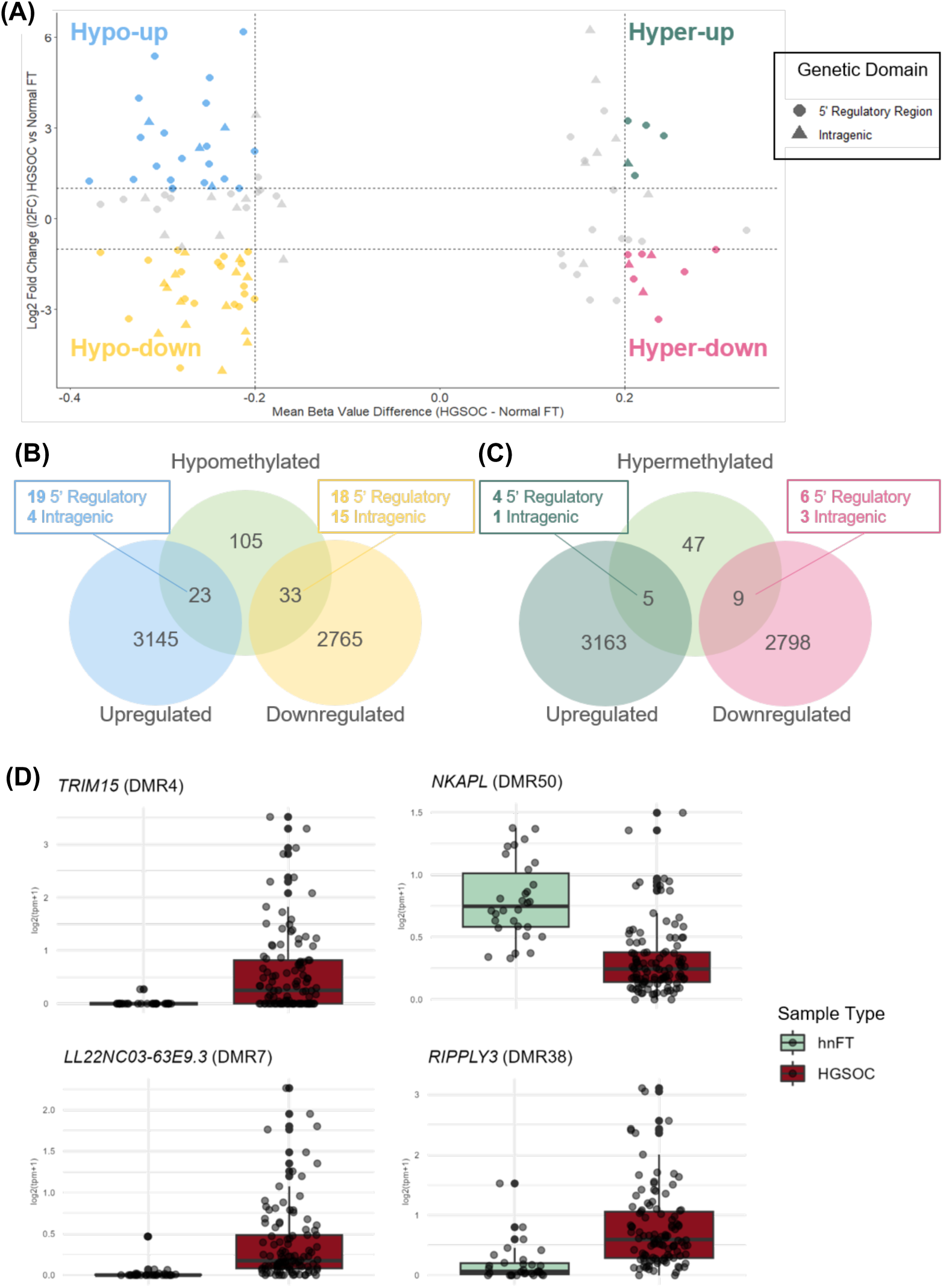
Integration of DNA methylation and an independent gene expression dataset. **(A)** Scatter plot of mean β value difference (HGSOC vs hnFT) vs log2 fold change (HGSOC vs hnFT). Each point represents a gene-DMR pair and point shape represents the gene region in which the DMR is methylated (5’ Regulatory region or intragenic). **(B)** Venn diagram summarising the intersection between hypomethylated genes and DEGs. (**C)** Venn diagram summarising the intersection between hypermethylated genes and DEGs. **(D**) Expression of the four DEGs (*TRIM15, LL22NC03-63E9.3, NKAPL* and *RIPPLY3*) located within the top 10 most significantly hypermethylated or hypomethylated DMRs.

Of the 161 genes located in hypomethylated DMRs, 23 were upregulated and 33 were downregulated (**Figure 5B**). Of the 61 genes located in hypermethylated DMRs, 5 were upregulated and 9 were downregulated (**Figure 5C**). Four differentially methylated and expressed genes (*TRIM15, LL22NC03-63E9.3, NKAPL and RIPPLY3*) were located within the top 10 most significantly hyper/hypomethylated regions (**Figure 4D**). *TRIM15* and *LL22NC03-63E9.3* are hypomethylated in their 5’ regulatory regions and exhibit elevated expression in HGSOC. *NKAPL* and *RIPPLY3* are hypermethylated within their 5’ regulatory regions, and *NKAPL* is downregulated whereas *RIPPLY3* is upregulated in HGSOC (**Figure 4D**). Gene Ontology analysis of inversely correlated DMRs showed involvement in pathway activation, for example in metabolic processes and gene expression (p value <0.01), (**Supplementary Figure 5).** Only two pathways were suppressed, specifically “organic cyclic compound binding” and “bounding membrane of organelle” (p value <0.04).

## Discussion

Mounting evidence supports epithelial cells from the fallopian tube, rather than the ovarian surface, as the origin of HGSOC^4^. Understanding the molecular and epigenetic changes that drive transformation from fallopian tube STIC to invasive carcinoma is therefore critical for uncovering early events in HGSOC development.

STICs carry HGSOC driver mutations, namely those in *TP53, BRCA1*, *BRCA2* or *PTEN,* suggesting that tumour identity is formed within these lesions^5^. One previous study demonstrated that focal DNA hypermethylation patterns of HGSOC also originate in STIC^9^. Here we report that the even more widespread global hypomethylation initiates in STICs, which becomes more extensive with progression to HGSOC. This observation is consistent with previous evidence that genome-wide DNA hypomethylation occurs early in tumorigenesis and increases with disease progression^28–30^. We observed similar quantities of DMCpGs and directional changes in HGSOC and STIC. However, STICs exhibited a higher frequency of hypermethylation than HGSOC, suggesting that specific hypermethylated regions may help maintain STIC identity, while their loss may mark progression to invasive HGSOC. STIC lesions are difficult to obtain for research purposes due to their small size; therefore, with a small dataset of only 11 STIC tissues, it is important that our findings are validated in future studies.

Notably, the differential DNA methylation common to STIC and HGSOC was nonrandomly distributed. As is typical, DNA hypermethylation was found predominantly at 5’ regulatory regions and CpG islands, whereas hypomethylation tended towards inter– and intragenic sequences and non-CpG dense open sea domains. In contrast to promoter hypermethylation, a surprising finding was the predominant hypomethylation of enhancer regulatory elements. Enhancer domains are known to be tissue specific, however cancer can hijack and alter the function of tissue specific enhancers^31^. We identified significant differences in DNA methylation between hnFT and STIC/HGSOC tissues across CpG sites within enhancers. However future functional studies are needed to confirm the impact of these alterations. There is also the debate as to which tissue is most relevant to study, to determine whether fallopian tube or ovarian tissue specific enhancers are more appropriate. This may reflect early activation of oncogenic transcriptional programs in STIC preceding HGSOC development and warrants further functional studies to elucidate the biological consequences of these epigenetic alterations. The DNA methylation of enhancers has previously been shown to be highly dynamic, supporting their reprogramming during cancer development and progression^31, 32^.

We also observed imbalances in DNA methylation across chromosomes, most notably hypomethylation of chromosome 18, and the complete absence of X chromosome hypermethylation, suggesting significant chromosomal instability in STIC and HGSOC. Chromosome 18 hypomethylation in HGSOC has not been previously reported, although q-arm deletions have been noted^33^. Chromosome 18 is particularly GC-poor, gene-sparse, and typically localised at the nuclear periphery within heterochromatin^34–36^. Beyond the scope of this study, future research should explore hypomethylation of chromosome 18 and its possible effect on nuclear positioning, transcriptional potential, and inter-chromosomal interactions. In healthy female cells, one X chromosome forms a Barr body, yet in some cancers, for example, breast, cervical and ovarian there has been a reported loss or absence of detectable Barr bodies^37, 38^ hypomethylation/reactivation of genes on the inactive X chromosome has been observed in HGSOC^39^. Future studies examining chromatin architecture in STIC would be beneficial in determining whether there is a loss of the X chromosome barr body prior to HGSOC onset.

The retention of 11,660 overlapping DMCpGs across independent DNA methylation datasets underscores the strong epigenetic continuity between STIC and HGSOC and supports the growing view in the field that HGSOC emerges from STIC precursors^40, 41^. The integration of RNA-sequencing data allowed us to identify genes differentially expressed between HGSOC and hnFT, revealing transcriptional changes associated with ovarian cancer initiation. A total of 5,966 DEGs were identified, with 3168 upregulated and 2798 downregulated, highlighting widespread transcriptional reprogramming in HGSOC. By intersecting these DEGs with genes mapped to DMRs, we identified 70 genes with concurrent differential methylation, suggesting a potential role of these genes in disease progression. We identified four genes (*TRIM15, LL22NC03-63E9.3, NKAPL*, and *RIPPLY3*) that were both differentially expressed and located within the top 10 most significant hyper/hypo methylated regions. Hypomethylation in the 5’ regulatory regions of both *TRIM15* and *LL22NC03-63E9.3* was observed in conjunction with overexpression in HGSOC, exemplifying how reduced methylation in regulatory regions may activate transcription. While *TRIM15*, the gene coding for a focal adhesion protein, has been associated with various cancer types, this is the first report of this gene’s potential involvement in HGSOC^42–44^. *LL22NC03-63E9.3*, an uncharacterized long non-coding RNA has also not been studied in the context of ovarian cancer. Both *NKAPL* and *RIPPLY3* were found to be hypermethylated in their 5’ regulator regions. We observed a downregulation of *NKAPL* in HGSOC, which aligns with the expected role of promoter methylation in repressing gene expression.

We previously reported *NKAPL* Hypermethylation in cfDNA as a prognostic predictor in HGSOC^45^. We showed that demethylation of the *NKAPL* promoter increased platinum sensitivity in cell line models, suggesting *NKAPL’s* potential role as a modulator of chemoresistance. Interestingly, while *RIPPLY3* was found to harbour hypermethylation in its 5’ regulatory region, we noted that this gene was in fact upregulated in HGSOC, which may suggest the involvement of additional regulatory mechanisms. Nonetheless, *RIPPLY3* has been studied as a blood-based biomarker for HGSOC^46^. One strength of this analysis is that the RNA-seq data was entirely independent of the methylation datasets, yet we still observed a significant overlap in genes that were both differentially expressed and methylated across HGSOC and hnFT tissues. However, it is important to note that these transcriptomic datasets represent a relatively small cohort of samples (n=24 HGSOC, n=27 hnFT). We unfortunately could not compare DNA methylation of STIC lesions due to the lack of publicly data available at the time of this analysis. These small samples sizes reduce statistical power, which limits the identification of more subtle transcriptional changes. Despite this, the strong overlap between DEGs and genes within DMRs indicates the biological relevance of our findings. While the use of an independent RNA-seq cohort provides a degree of external validation, the absence of matched methylation and transcriptomic profiling from the same samples means that direct, sample-level correlations between DMR methylation status and gene expression cannot be established. Interpatient tumour heterogeneity, a well-recognised feature of HGSOC, may further confound cross-dataset comparisons, as differences in tumour microenvironment, clonal composition, and stromal admixture may differentially influence methylation and expression profiles across cohorts. Although batch effects between datasets were accounted for statistically where possible, residual technical variation inherent to independently generated datasets cannot be fully excluded. Future studies employing matched, multi-omic profiling within the same patient cohort would be necessary to definitively establish causal relationships between the methylation changes identified here and their transcriptional consequences.

There is a challenge associated with selecting appropriate control samples when studying ovarian cancer. Acquiring histologically normal ovarian epithelium is extremely difficult, and subsequently not often used. Indeed, no EPIC array data are available for normal ovarian epithelium. Fallopian tube tends to be used, and it is easily obtained during surgery. In addition, molecular analysis has shown more similarities between HGSOC and hnFT, than healthy ovarian epithelium or peritoneum^47^, supporting the use of hnFT as a suitable control. These are obviously also very appropriate for studying STIC lesions. Limited quantities of dSTIC and p53 lesion quantities limited our ability to fully understand their role in disease progression and STIC onset. Obtaining sufficient sample sizes for these tissues is notoriously problematic, owing largely to difficulty in acquiring such samples due to rarity or microscopic size, resulting in them being consumed in routine histological analysis. Studies, such as this one, albeit small, are thus extremely important. We show that a dSTIC lesion demonstrates DNA methylation patterns similar to STIC, with a p53 lesion demonstrating similarities to hnFT. If larger cohorts could be obtained, future research could compare the DNA methylation of p53 lesions, serous tubal intraepithelial lesions (STILs)/dSTICs with proliferatively active STICs. STIC tissues were obtained via laser-capture microdissection, thereby enhancing sample purity. However, tumour content could not be confirmed for the HGSOC tissues, highlighting a recurrent challenge associated with publicly available datasets and the constraints of the metadata provided.

Our findings reveal that epigenetic reprogramming, particularly altered DNA methylation, occurs before the onset of HGSOC and likely underpins its initiation and progression. Specifically, canonical cancer methylation hallmarks, regional hypermethylation of functional elements alongside global hypomethylation, emerge first in STICs and intensify with disease evolution, suggesting a driving role in early transformation and metastatic spread. The striking overlap in methylation signatures between STIC and HGSOC supports the concept that STICs represent a molecularly advanced, pre-invasive stage of disease. Comprehensive validation of these early epigenetic alterations will be critical for developing biomarkers of early detection and uncovering mechanisms of chromatin deregulation that enable metastasis. Addressing the current gap in early-stage HGSOC research by integrating epigenetic profiling of precursor lesions like STICs offers a powerful route to intercept ovarian cancer at its molecular origin.

## Acknowledgements

We would like to thank the Irish Research Council and Breakthrough Cancer Research for funding this project through the Enterprise Partnership Scheme (EPSPG/2021/169). We would also like to thank Dr Kim Conteddu and Unmani Jaygude for their bioinformatic and statistical support.

## SUPPLEMENTARY TABLES

**Supplementary Table 1.**
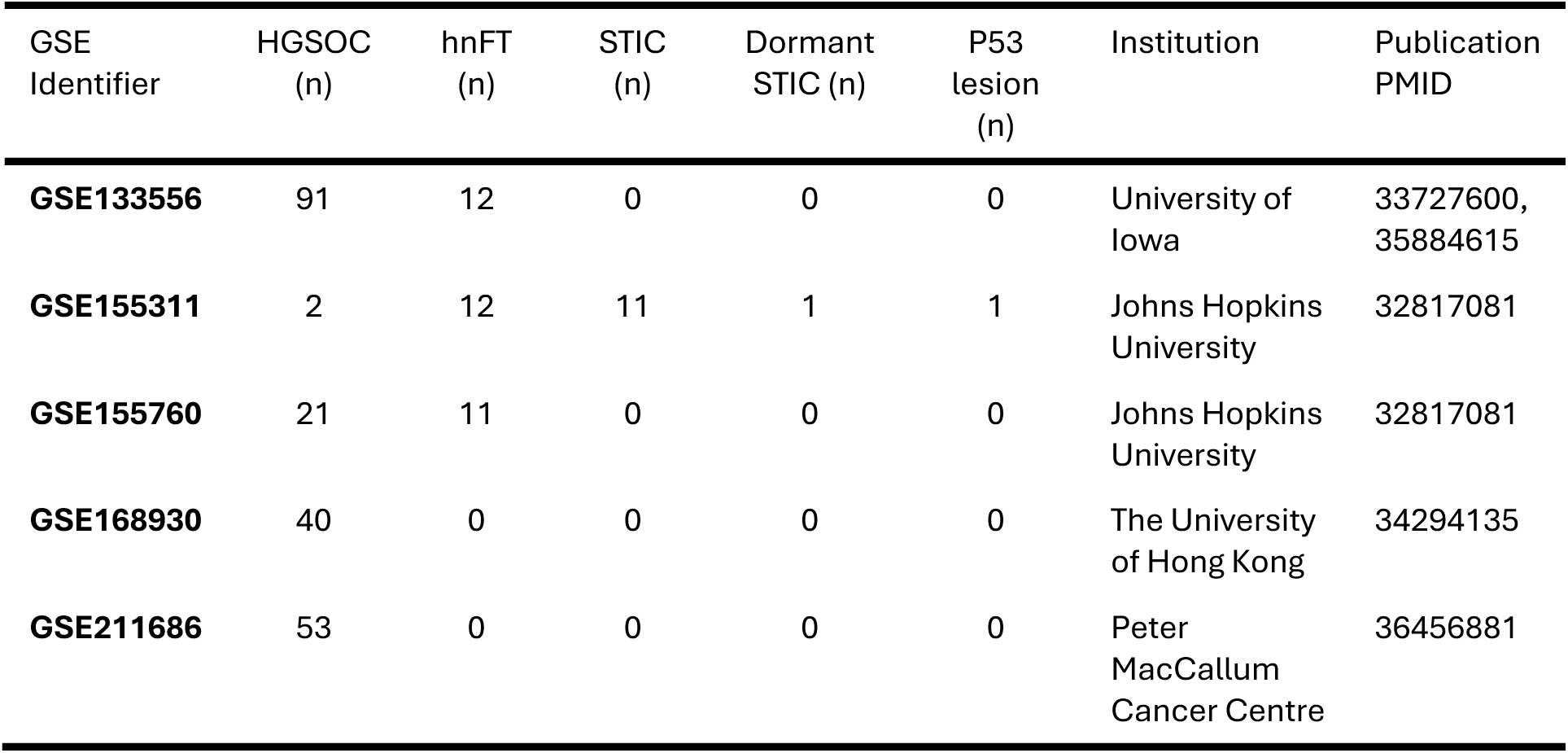
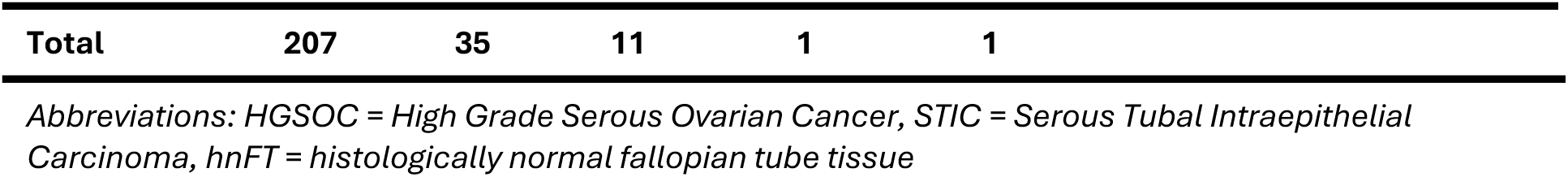
Source of datasets and samples.

| GSE Identifier | HGSOC (n) | hnFT (n) | STIC (n) | Dormant STIC (n) | P53 lesion (n) | Institution | Publication PMID |
| --- | --- | --- | --- | --- | --- | --- | --- |
| **GSE133556** | 91 | 12 | 0 | 0 | 0 | University of Iowa | 33727600, 35884615 |
| **GSE155311** | 2 | 12 | 11 | 1 | 1 | Johns Hopkins University | 32817081 |
| **GSE155760** | 21 | 11 | 0 | 0 | 0 | Johns Hopkins University | 32817081 |
| **GSE168930** | 40 | 0 | 0 | 0 | 0 | The University of Hong Kong | 34294135 |
| **GSE211686** | 53 | 0 | 0 | 0 | 0 | Peter MacCallum Cancer Centre | 36456881 |

|  |  |  |  |  |  |
| --- | --- | --- | --- | --- | --- |
| **Total** | **207** | **35** | **11** | **1** | **1** |

Abbreviations: HGSOC = High Grade Serous Ovarian Cancer, STIC = Serous Tubal Intraepithelial Carcinoma, hnFT = histologically normal fallopian tube tissue

**Supplementary Table 2.** The quantities of probes filtered during data processing.

| Cohort | Probes that cover SNPs within their last 5 bases (n) | Cross reactive probes (n) | Poor performing probes (n) | Poor performing samples (n) | Non-CpG probes (n) | Total probes removed (n) | Total probes remaining (n) |
| --- | --- | --- | --- | --- | --- | --- | --- |
| **Unmatched** | 21,414 | 43,140 | 22,261 | 4 | 1,178 | 87,993 | **778,902** |
| **Matched** | 21,414 | 43,140 | 37,243 | 0 | 951 | 102,748 | **764,147** |
| **Verification** | 21,414 | 43,140 | 8,686 | 0 | 1,315 | 74,555 | **792,340** |

**Supplementary Table 3.** Differentially Methylated Regions (DMRs) in HGSOC.

|  | All DMRs | Hypermethylated DMRs | Hypomethylated DMRs |
| --- | --- | --- | --- |
| **Total DMRs n, %** | **N (%)** |  | **342, 76.51%** |
| Total | 447 (100) | **105 (23.49)** |  |
| Number of significant DMCPGs per DMR, mean | 3.7 | 4.0 | 3.6 |
| Number of significant DMCPGs per DMR, range | 3 - 12 | 3 - 12 | 3 - 10 |
| Mean DMR length (base pairs) | 573.3 | 604.6 | 563.7 |
| Length range (min-max) | 7 – 2,916 | 12 – 2,591 | 7 – 2,916 |
| Mean β-value | -0.15 | 0.19 | -0.26 |
| Mean β-value range (min-max) | 0.13 - 0.40 | 0.13 - 0.33 | 0.15 - 0.40 |

## SUPPLEMENTARY FIGURES

**Supplementary Figure 1.**
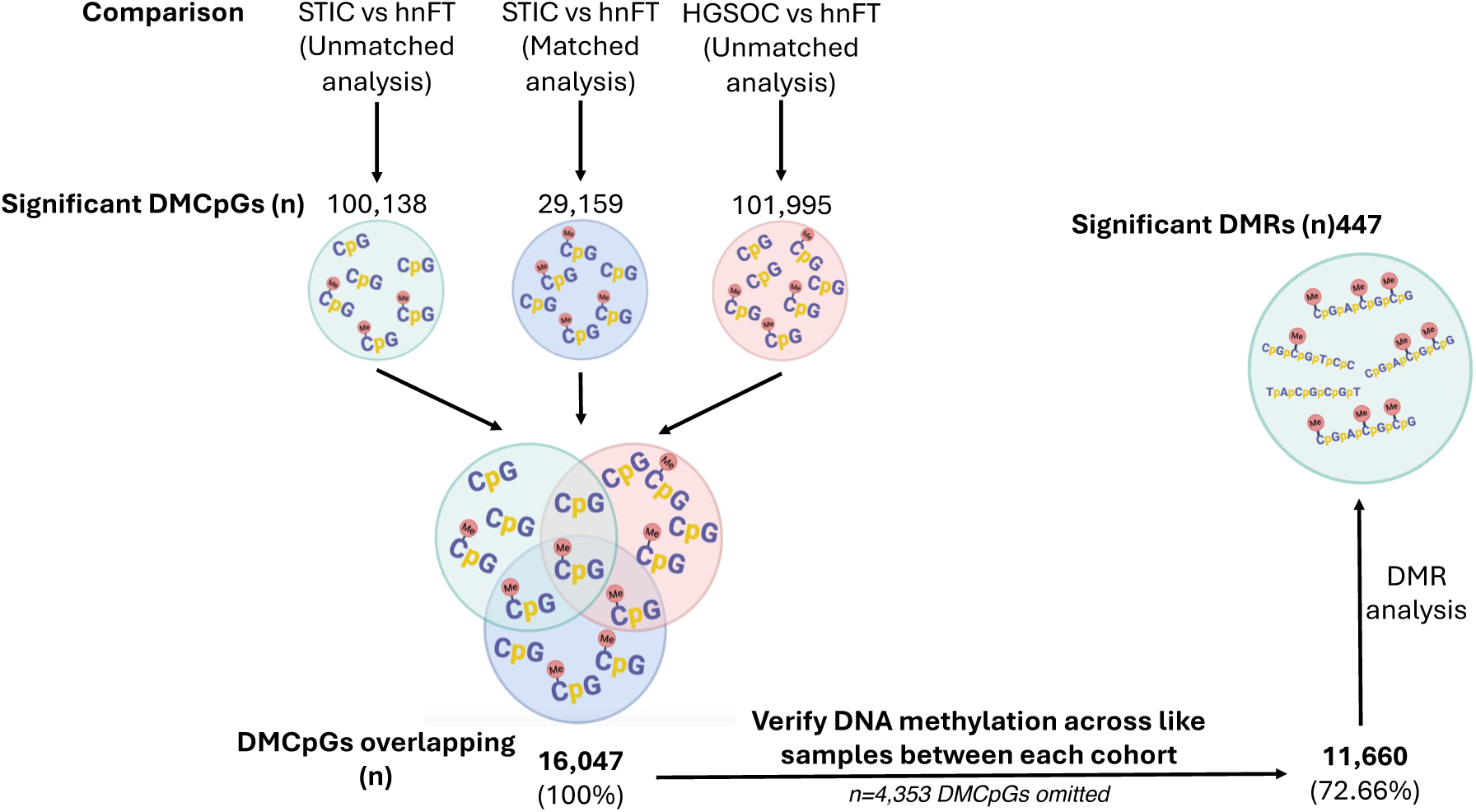
Flow diagram detailing the process of identifying overlapping CpG sites.

**Supplementary Figure 2.**
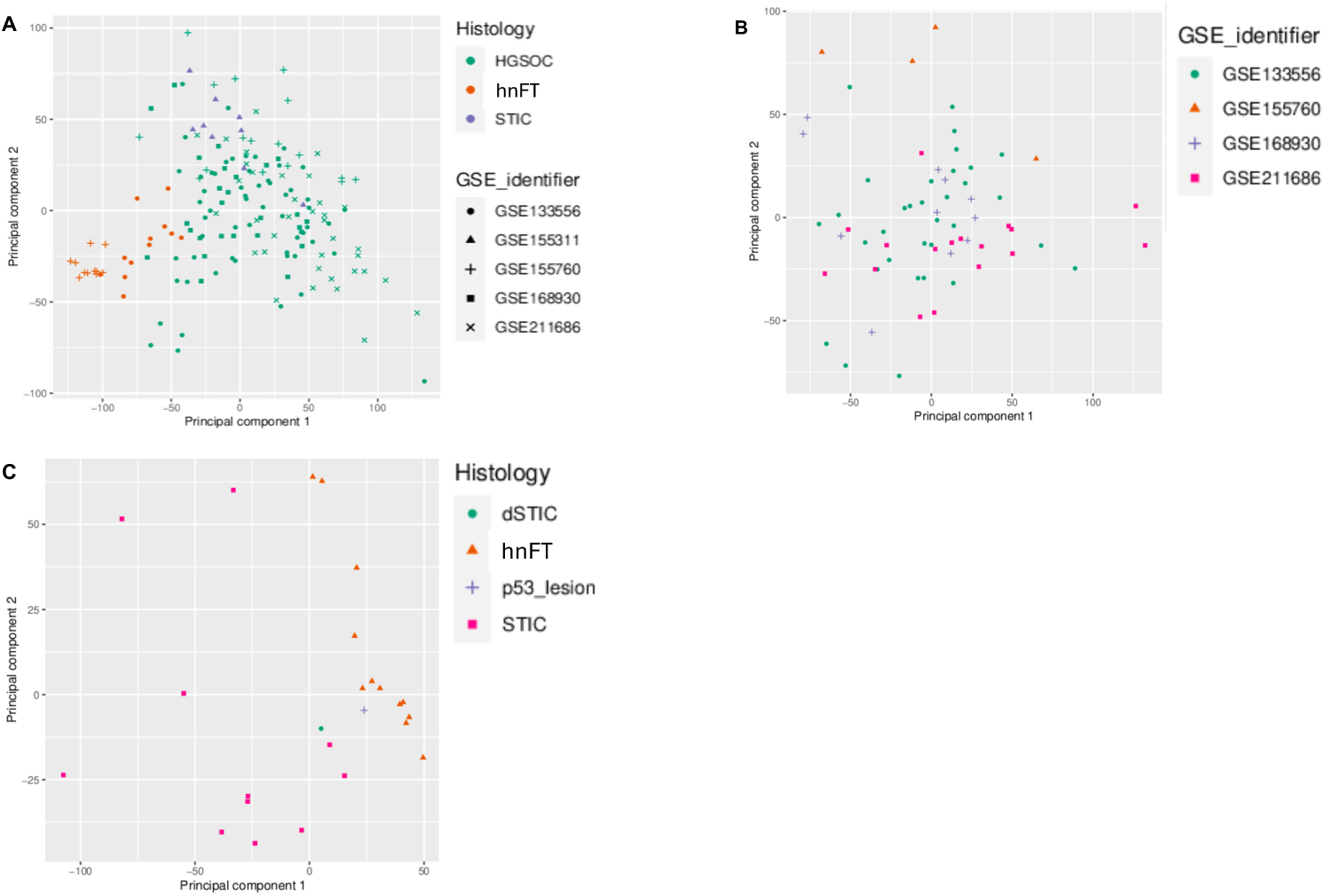
Principal component analysis of EPIC DNA methylation array data. (**A**) discovery cohort, (**B**) validation cohort (HGSOC tissues only) and (**C**) matched cohort.

**Supplementary Figure 3.**
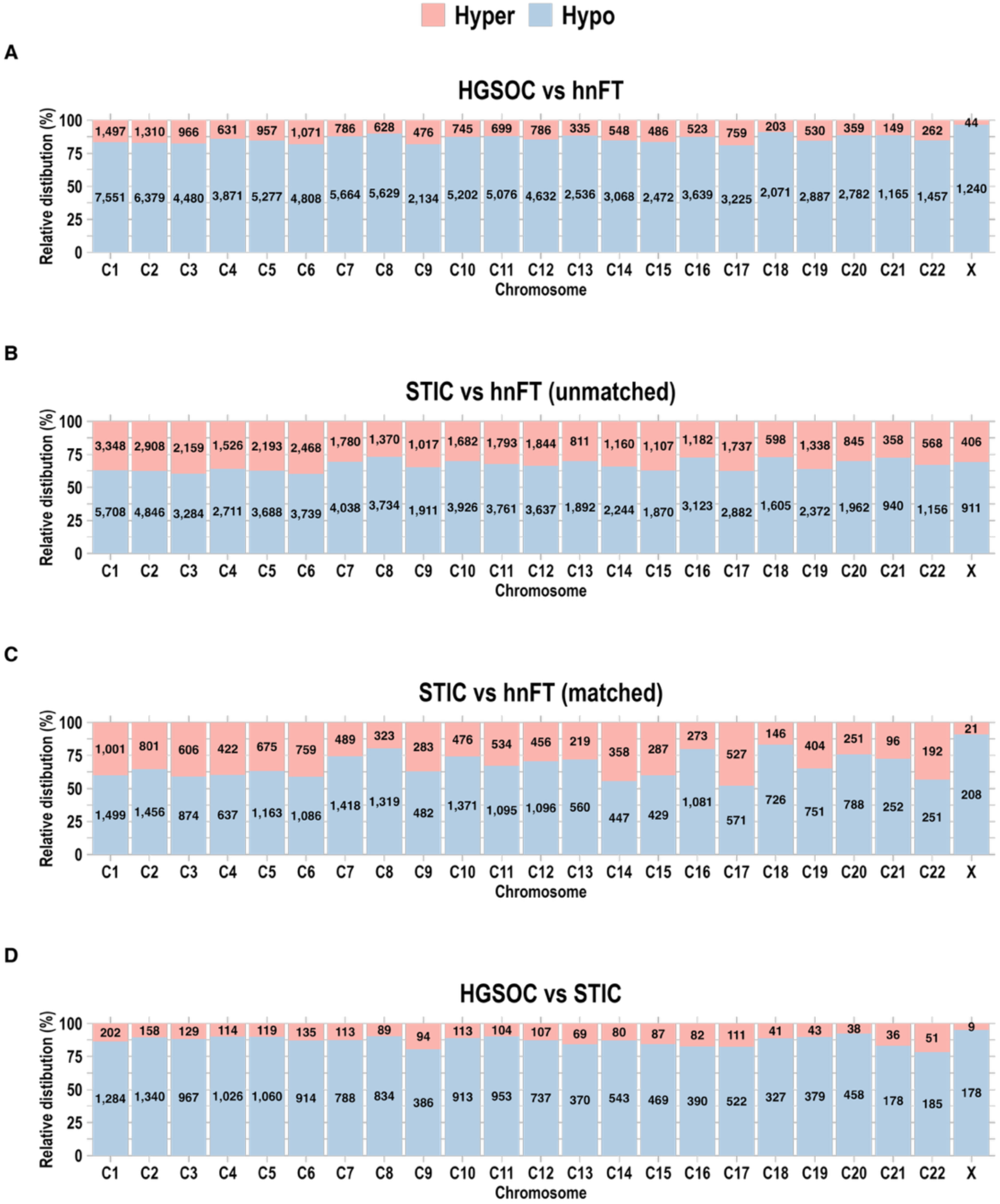
Chromosomal annotation of DMCpGs between all comparisons. Stacked bar charts indicating the adjusted percentage of the hypermethylated (Hyper) and hypomethylated (Hypo) DMCpGs chromosomal distributions for each comparison (**A**) HGSOC vs hnFT (unmatched) (**B**) STIC vs hnFT (unmatched), (**C**) STIC vs hnFT (matched) (**D**) HGSOC vs STIC (unmatched). from unmatched STIC and hnFT tissues (i.e. tissues derived from different patients). The y-axis shows the relative distribution (%) of each CpG site on each chromosome. A chi-squared test of independence comparing proportions of hypermethylated and hypomethylated sites across chromosomes yielded a statistically significant p-value < 0.0001).

**Supplementary Figure 4.**
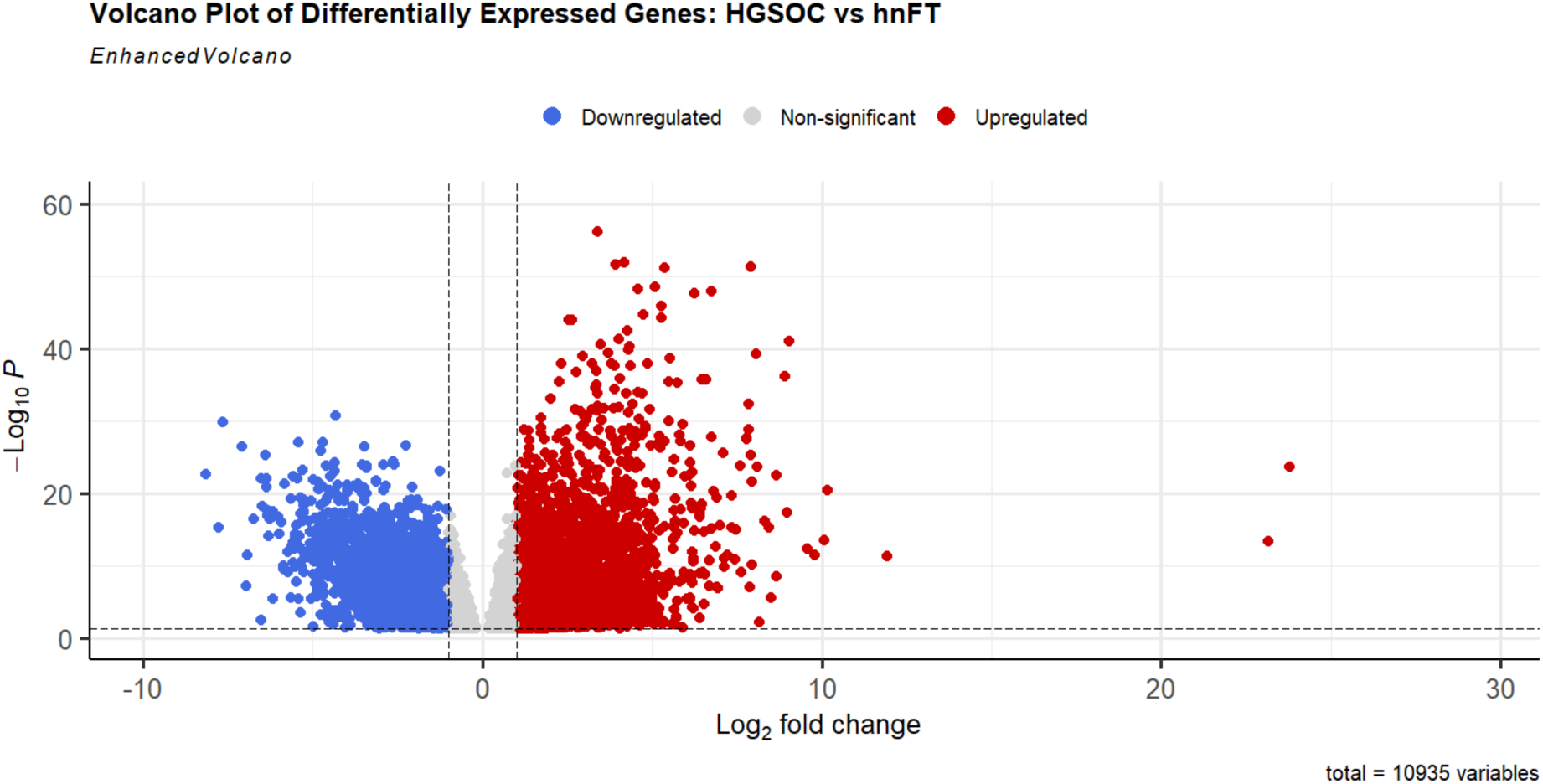
Gene deregulation in HGSOC vs hnFT samples. The x-axis represents the log2 fold change (L2FC) in gene expression, with positive values indicating upregulation in HGSOC and negative values indicating downregulation. The y-axis represents the –log10 of the adjusted p-value, indicating statistical significance. Genes with a log2 fold change greater than 1 (or less than –1) and an adjusted p-value < 0.05 are considered significantly differentially expressed and are highlighted in red (upregulated) and blue (downregulated). Non-significant genes are shown in grey.

**Supplementary Figure 5.**
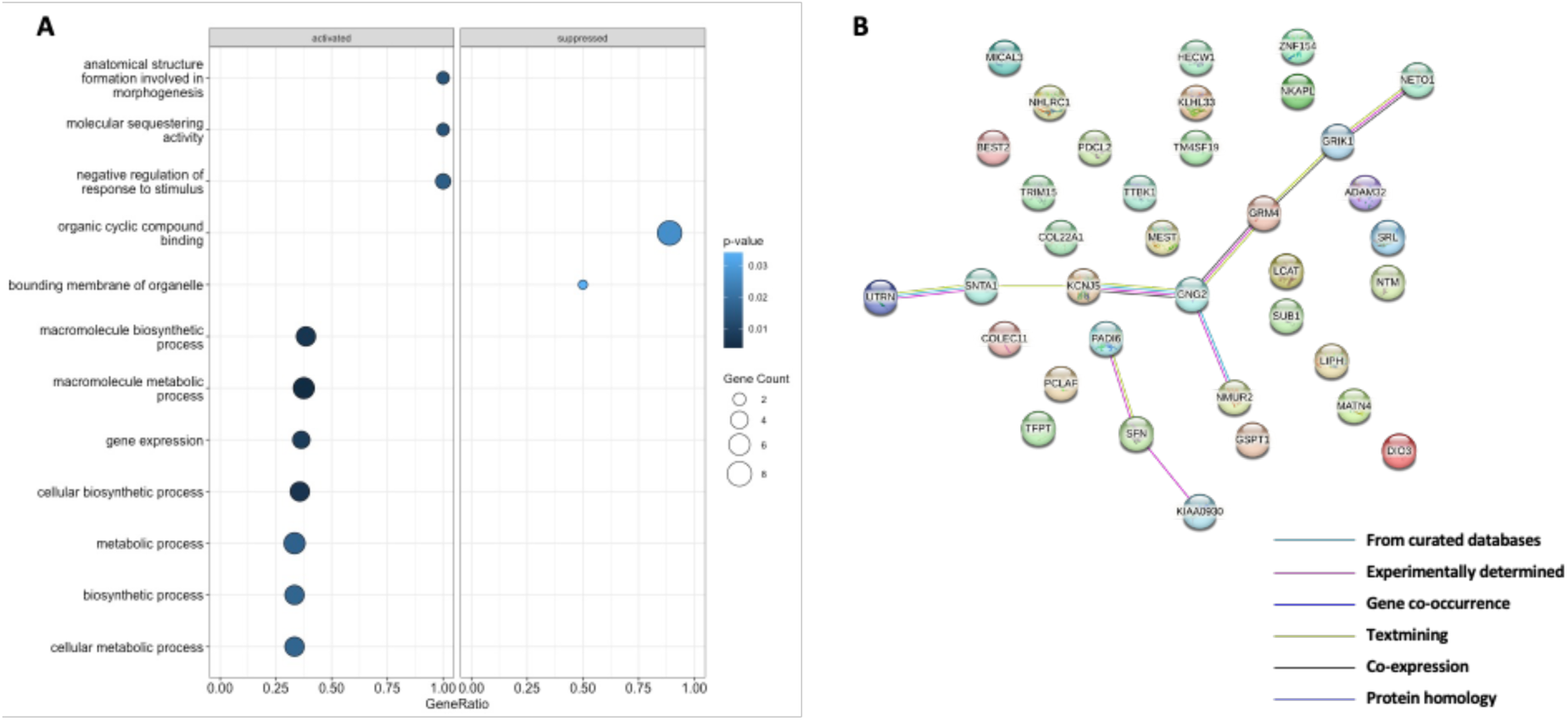
Gene Ontology (GO) Enrichment and protein-protein interaction (PPI) analysis on inversely correlated DMRs and associated genes. **(A)** GO enrichment analysis of genes associated with inversely correlated DMRs. This is shown for both activated and suppressed gene sets based on gene expression ranking. Dot plot is coloured according to non-adjusted p-value. The X-axis represents the Gene Ratio, defined as the ratio of core enrichment genes to the total number of genes associated with each GO term (listed on Y-axis). Dot size corresponds to the number of genes (Count). **(B)** STRING PPI analysis of proteins associated with inversely correlated DMRs shows predicted and known interactions among differentially enriched genes based on STRING database analysis. Each node represents a protein, and edges indicate protein–protein associations derived from curated databases, experimental data, co-expression, gene neighbourhood, and text mining. Thicker or coloured edges denote stronger or multiple lines of evidence for interaction. Unconnected nodes are displayed to preserve the biological context of the network.

## References

1. American Cancer Society. Cancer Facts & Figures 2025: American Cancer Society, 2025.

2. Koshiyama M, Matsumura N, Konishi I. Subtypes of Ovarian Cancer and Ovarian Cancer Screening. Diagnostics (Basel) 2017;7.

3. Punzón-Jiménez P, Lago V, Domingo S, Simón C, Mas A. Molecular Management of High-Grade Serous Ovarian Carcinoma. International journal of molecular sciences 2022;23.

4. Colvin EK, Howell VM. Why the dual origins of high grade serous ovarian cancer matter 2020;11: 1200.

5. Labidi-Galy SI, Papp E, Hallberg D, Niknafs N, Adleff V, Noe M, Bhattacharya R, Novak M, Jones S, Phallen J, Hruban CA, Hirsch MS, Lin DI, Schwartz L, Maire CL, Tille J-C, Bowden M, Ayhan A, Wood LD, Scharpf RB, Kurman R, Wang T-L, Shih I-M, Karchin R, Drapkin R, Velculescu VE. High grade serous ovarian carcinomas originate in the fallopian tube 2017;8: 1093.

6. Bogaerts JMA, Steenbeek MP, van Bommel MHD, Bulten J, van der Laak JAWM, de Hullu JA, Simons M. Recommendations for diagnosing STIC: a systematic review and meta-analysis 2022;480: 725–37.

7. Locke WJ, Guanzon D, Ma C, Liew YJ, Duesing KR, Fung KYC, Ross JP. DNA Methylation Cancer Biomarkers: Translation to the Clinic. Front Genet 2019;10: 1150.

8. Chan DW, Lam WY, Chen F, Yung MMH, Chan YS, Chan WS, He F, Liu SS, Chan KKL, Li B, Ngan HYS. Genome-wide DNA methylome analysis identifies methylation signatures associated with survival and drug resistance of ovarian cancers. Clin Epigenetics 2021;13: 142.

9. Pisanic TR, 2nd, Wang Y, Sun H, Considine M, Li L, Wang TH, Wang TL, Shih IM. Methylomic Landscapes of Ovarian Cancer Precursor Lesions. Clin Cancer Res 2020;26: 6310–20.

10. Cardillo N, Devor EJ, Pedra Nobre S, Newtson A, Leslie K, Bender DP, Smith BJ, Goodheart MJ, Gonzalez-Bosquet J. Integrated Clinical and Genomic Models to Predict Optimal Cytoreduction in High-Grade Serous Ovarian Cancer. Cancers (Basel) 2022;14.

11. Gonzalez Bosquet J, Devor EJ, Newtson AM, Smith BJ, Bender DP, Goodheart MJ, McDonald ME, Braun TA, Thiel KW, Leslie KK. Creation and validation of models to predict response to primary treatment in serous ovarian cancer. Scientific reports 2021;11: 5957.

12. Garsed DW, Pandey A, Fereday S, Kennedy CJ, Takahashi K, Alsop K, Hamilton PT, Hendley J, Chiew YE, Traficante N, Provan P, Ariyaratne D, Au-Yeung G, Bateman NW, Bowes L, Brand A, Christie EL, Cunningham JM, Friedlander M, Grout B, Harnett P, Hung J, McCauley B, McNally O, Piskorz AM, Saner FAM, Vierkant RA, Wang C, Winham SJ, Pharoah PDP, Brenton JD, Conrads TP, Maxwell GL, Ramus SJ, Pearce CL, Pike MC, Nelson BH, Goode EL, DeFazio A, Bowtell DDL. The genomic and immune landscape of long-term survivors of high-grade serous ovarian cancer. Nat Genet 2022;54: 1853–64.

13. Saner FAM, Takahashi K, Budden T, Pandey A, Ariyaratne D, Zwimpfer TA, Meagher NS, Fereday S, Twomey L, Pishas KI, Hoang T, Bolithon A, Traficante N, Alsop K, Christie EL, Kang EY, Nelson GS, Ghatage P, Lee CH, Riggan MJ, Alsop J, Beckmann MW, Boros J, Brand AH, Brooks-Wilson A, Carney ME, Coulson P, Courtney-Brooks M, Cushing-Haugen KL, Cybulski C, El-Bahrawy MA, Elishaev E, Erber R, Gayther SA, Gentry-Maharaj A, Gilks CB, Harnett PR, Harris HR, Hartmann A, Hein A, Hendley J, Hernandez BY, Jakubowska A, Jimenez-Linan M, Jones ME, Kaufmann SH, Kennedy CJ, Kluz T, Koziak JM, Kristjansdottir B, Le ND, Lener M, Lester J, Lubiński J, Mateoiu C, Orsulic S, Ruebner M, Schoemaker MJ, Shah M, Sharma R, Sherman ME, Shvetsov YB, Soong TR, Steed H, Sukumvanich P, Talhouk A, Taylor SE, Vierkant RA, Wang C, Widschwendter M, Wilkens LR, Winham SJ, Anglesio MS, Berchuck A, Brenton JD, Campbell I, Cook LS, Doherty JA, Fasching PA, Fortner RT, Goodman MT, Gronwald J, Huntsman DG, Karlan BY, Kelemen LE, Menon U, Modugno F, Pharoah PDP, Schildkraut JM, Sundfeldt K, Swerdlow AJ, Goode EL, DeFazio A, Köbel M, Ramus SJ, Bowtell DDL, Garsed DW. Concurrent RB1 Loss and BRCA Deficiency Predicts Enhanced Immunologic Response and Long-term Survival in Tubo-ovarian High-grade Serous Carcinoma. Clin Cancer Res 2024;30: 3481–98.

14. Barrett T, Wilhite SE, Ledoux P, Evangelista C, Kim IF, Tomashevsky M, Marshall KA, Phillippy KH, Sherman PM, Holko M, Yefanov A, Lee H, Zhang N, Robertson CL, Serova N, Davis S, Soboleva A. NCBI GEO: archive for functional genomics data sets--update. Nucleic acids research 2013;41: D991–5.

15. Müller F, Scherer M, Assenov Y, Lutsik P, Walter J, Lengauer T, Bock C. RnBeads 2.0: comprehensive analysis of DNA methylation data 2019;20: 55.

16. Maksimovic J, Gordon L, Oshlack A. SWAN: Subset-quantile Within Array Normalization for Illumina Infinium HumanMethylation450 BeadChips 2012;13: R44.

17. Leek JT, Johnson WE, Parker HS, Jaffe AE, Storey JD. The sva package for removing batch effects and other unwanted variation in high-throughput experiments. Bioinformatics 2012;28: 882–3.

18. Gu Z, Eils R, Schlesner M. Complex heatmaps reveal patterns and correlations in multidimensional genomic data. Bioinformatics 2016;32: 2847–9.

19. Pidsley R, Zotenko E, Peters TJ, Lawrence MG, Risbridger GP, Molloy P, Van Djik S, Muhlhausler B, Stirzaker C, Clark SJ. Critical evaluation of the Illumina MethylationEPIC BeadChip microarray for whole-genome DNA methylation profiling. Genome biology 2016;17: 208.

20. Peters TJ, Meyer B, Ryan L, Achinger-Kawecka J, Song J, Campbell EM, Qu W, Nair S, Loi-Luu P, Stricker P, Lim E, Stirzaker C, Clark SJ, Pidsley R. Characterisation and reproducibility of the HumanMethylationEPIC v2.0 BeadChip for DNA methylation profiling 2024;25: 251.

21. Ritchie ME, Phipson B, Wu D, Hu Y, Law CW, Shi W, Smyth GK. limma powers differential expression analyses for RNA-sequencing and microarray studies 2015;43: e47-e.

22. O’Cathail C, Ahamed A, Burgin J, Cummins C, Devaraj R, Gueye K, Gupta D, Gupta V, Haseeb M, Ihsan M, Ivanov E, Jayathilaka S, Kadhirvelu V, Kumar M, Lathi A, Leinonen R, McKinnon J, Meszaros L, Pauperio J, Pesant S, Rahman N, Rinck G, Selvakumar S, Suman S, Sunthornyotin Y, Ventouratou M, Waheed Z, Woollard P, Yuan D, Zyoud A, Burdett T, Cochrane G. The European Nucleotide Archive in 2024 2024: gkae975.

23. Andrews S. FastQC: a quality control tool for high throughput sequence data., 2010.

24. Krueger F. Trim galore. A wrapper tool around Cutadapt and FastQC to consistently apply quality and adapter trimming to FastQ files, 2015.

25. Patro R, Duggal G, Love MI, Irizarry RA, Kingsford C. Salmon provides fast and bias-aware quantification of transcript expression 2017;14: 417–9.

26. Love MI, Huber W, Anders S. Moderated estimation of fold change and dispersion for RNA-seq data with DESeq2 2014;15: 550.

27. Skvortsova K, Stirzaker C, Taberlay P. The DNA methylation landscape in cancer 2019;63: 797–811.

28. Ehrlich M. DNA hypomethylation in cancer cells. Epigenomics 2009;1: 239–59.

29. Keita M, Wang Z-Q, Pelletier J-F, Bachvarova M, Plante M, Gregoire J, Renaud M-C, Mes-Masson A-M, Paquet ÉR, Bachvarov D. Global methylation profiling in serous ovarian cancer is indicative for distinct aberrant DNA methylation signatures associated with tumor aggressiveness and disease progression 2013;128: 356–63.

30. Earp MA, Cunningham JM. DNA methylation changes in epithelial ovarian cancer histotypes 2015;106: 311–21.

31. Yang J, Zhou F, Luo X, Fang Y, Wang X, Liu X, Xiao R, Jiang D, Tang Y, Yang G, You L, Zhao Y. Enhancer reprogramming: critical roles in cancer and promising therapeutic strategies 2025;11: 84.

32. Ziller MJ, Gu H, Müller F, Donaghey J, Tsai LTY, Kohlbacher O, De Jager PL, Rosen ED, Bennett DA, Bernstein BE, Gnirke A, Meissner A. Charting a dynamic DNA methylation landscape of the human genome 2013;500: 477–81.

33. Arnold N, Hägele L, Walz L, Schempp W, Pfisterer J, Bauknecht T, Kiechle M. Overrepresentation of 3q and 8q material and loss of 18q material are recurrent findings in advanced human ovarian cancer. *Genes*, Chromosomes and Cancer 1996;16: 46–54.

34. Nusbaum C, Zody MC, Borowsky ML, Kamal M, Kodira CD, Taylor TD, Whittaker CA, Chang JL, Cuomo CA, Dewar K, FitzGerald MG, Yang X, Abouelleil A, Allen NR, Anderson S, Bloom T, Bugalter B, Butler J, Cook A, DeCaprio D, Engels R, Garber M, Gnirke A, Hafez N, Hall JL, Norman CH, Itoh T, Jaffe DB, Kuroki Y, Lehoczky J, Lui A, Macdonald P, Mauceli E, Mikkelsen TS, Naylor JW, Nicol R, Nguyen C, Noguchi H, O’Leary SB, O’Neill K, Piqani B, Smith CL, Talamas JA, Topham K, Totoki Y, Toyoda A, Wain HM, Young SK, Zeng Q, Zimmer AR, Fujiyama A, Hattori M, Birren BW, Sakaki Y, Lander ES. DNA sequence and analysis of human chromosome 18. Nature 2005;437: 551–5.

35. Federico C, Bruno F, Ragusa D, Clements CS, Brancato D, Henry MP, Bridger JM, Tosi S, Saccone S. Chromosomal Rearrangements and Altered Nuclear Organization: Recent Mechanistic Models in Cancer. Cancers (Basel) 2021;13.

36. Croft JA, Bridger JM, Boyle S, Perry P, Teague P, Bickmore WA. Differences in the localization and morphology of chromosomes in the human nucleus. J Cell Biol 1999;145: 1119–31.

37. Pageau GJ, Hall LL, Ganesan S, Livingston DM, Lawrence JB. The disappearing Barr body in breast and ovarian cancers Nat Rev Cancered., vol. 7. England, 2007: 628–33.

38. Kristiansen M, Helland A, Kristensen GB, Olsen AO, Lønning PE, Børresen-Dale AL, Ørstavik KH. X chromosome inactivation in cervical cancer patients. Cancer Genet Cytogenet 2003;146: 73–6.

39. Kang J, Lee HJ, Kim J, Lee JJ, Maeng LS. Dysregulation of X chromosome inactivation in high grade ovarian serous adenocarcinoma. PLoS One 2015;10: e0118927.

40. Pisanic TR, 2nd, Cope LM, Lin SF, Yen TT, Athamanolap P, Asaka R, Nakayama K, Fader AN, Wang TH, Shih IM, Wang TL. Methylomic Analysis of Ovarian Cancers Identifies Tumor-Specific Alterations Readily Detectable in Early Precursor Lesions. Clin Cancer Res 2018;24: 6536–47.

41. Giancontieri P, Turetta C, Barchiesi G, Pernazza A, Pignataro G, D’Onghia G, Santini D, Tomao F. High-grade serous carcinoma of unknown primary origin associated with STIC clinically presented as isolated inguinal lymphadenopathy: a case report. Front Oncol 2023;13: 1307573.

42. Lee O-H, Lee J, Lee KH, Woo YM, Kang J-H, Yoon H-G, Bae S-K, Songyang Z, Oh SH, Choi Y. Role of the focal adhesion protein TRIM15 in colon cancer development 2015;1853: 409–21.

43. Chen W, Lu C, Hong J. TRIM15 Exerts Anti-Tumor Effects Through Suppressing Cancer Cell Invasion in Gastric Adenocarcinoma. Med Sci Monit 2018;24: 8033–41.

44. Sun Y, Ren D, Yang C, Yang W, Zhao J, Zhou Y, Jin X, Wu H. TRIM15 promotes the invasion and metastasis of pancreatic cancer cells by mediating APOA1 ubiquitination and degradation 2021;1867: 166213.

45. Silva R, Glennon K, Metoudi M, Moran B, Salta S, Slattery K, Treacy A, Martin T, Shaw J, Doran P, Lynch L, Jeronimo C, Perry AS, Brennan DJ. Unveiling the epigenomic mechanisms of acquired platinum-resistance in high-grade serous ovarian cancer. Int J Cancer 2023;153: 120–32.

46. Marinelli LM, Kisiel JB, Slettedahl SW, Mahoney DW, Lemens MA, Shridhar V, Taylor WR, Staub JK, Cao X, Foote PH, Burger KN, Berger CK, O’Connell MC, Doering KA, Giakoumopoulos M, Berg H, Volkmann C, Solsrud A, Allawi HT, Kaiser M, Vaccaro AM, Albright Crawford C, Moehlenkamp C, Shea G, Deist MS, Schoolmeester JK, Kerr SE, Sherman ME, Bakkum-Gamez JN. Methylated DNA markers for plasma detection of ovarian cancer: Discovery, validation, and clinical feasibility. Gynecol Oncol 2022;165: 568–76.

47. Ducie J, Dao F, Considine M, Olvera N, Shaw PA, Kurman RJ, Shih IM, Soslow RA, Cope L, Levine DA. Molecular analysis of high-grade serous ovarian carcinoma with and without associated serous tubal intra-epithelial carcinoma. Nature communications 2017;8: 990.

